# Trk Agonist Drugs Rescue Noise-Induced Hidden Hearing Loss

**DOI:** 10.1101/2020.07.01.182931

**Authors:** Katharine A. Fernandez, Takahisa Watabe, Mingjie Tong, Xiankai Meng, Kohsuke Tani, Sharon G. Kujawa, Albert S. B. Edge

## Abstract

TrkB agonist drugs are shown here to have a significant effect on the regeneration of afferent cochlear synapses after noise-induced synaptopathy. The effects were consistent with regeneration of cochlear synapses that we observed *in vitro* after synaptic loss due to kainic acid-induced glutamate toxicity and were elicited by administration of TrkB agonists, amitriptyline and 7,8-dihydroxyflavone, directly into the cochlea via the posterior semicircular canal 48 h after exposure to noise. Synaptic counts at the inner hair cell and wave 1 amplitudes in the ABR were partially restored 2 weeks after drug treatment. Effects of amitriptyline on wave 1 amplitude and afferent auditory synapse numbers in noise-exposed ears after systemic (as opposed to local) delivery were profound and long-lasting; synapses in the treated animals remained intact one year after the treatment. However, the effect of systemically delivered amitriptyline on synaptic rescue was dependent on dose and the time window of administration: it was only effective when given before noise exposure at the highest injected dose. The long-lasting effect and the efficacy of post-exposure treatment indicate a potential broad application for the treatment of synaptopathy, which often goes undetected until well after the original damaging exposure(s).

## Introduction

Sensorineural hearing loss results from pathology of the inner ear and its primary cause can reside in sensory hair cells or afferent neurons (Kujawa and Liberman, 2006, 2009; Nadol, 1997; Ruel et al., 2007; Sergeyenko et al., 2013; Wang et al., 2002; Wong and Ryan, 2015). However, work in animal models of noise- and age-related hearing loss has revealed that loss of synapses between the two is commonly the earliest finding (Kujawa and Liberman, 2009). This type of damage has been referred to as ‘hidden hearing loss’ because its presence is not captured by the threshold audiogram, the standard clinical assay of hearing loss. Although synaptic loss is not revealed by threshold measures until near total, it can be detected as a decline in the suprathreshold amplitudes of neural responses (Fernandez et al., 2020; Kujawa and Liberman, 2009; Sergeyenko et al., 2013).

Neurotrophins have been tested for activity in ameliorating noise- and toxin-induced damage to the neurons of the inner ear (Ramekers et al., 2015; Ruel et al., 2007; Shepherd et al., 2005; Shinohara et al., 2002; Wise et al., 2005). These molecules have numerous functions in neurons, including support for neural survival, outgrowth of neurites, and both guidance and synapse formation. While neurotrophins have been extensively tested as therapeutic agents, the difficulty in targeting therapeutic levels of proteins to the brain or peripheral nerves has hampered their use (Poduslo and Curran, 1996). In contrast, small molecules can reach target tissues in the nervous system, with demonstrated protective and regenerative effects (Jang et al., 2009).

Here, we examined the potential of amitriptyline (AT) and 7,8-dihydroxyflavone (DHF), structurally unrelated small molecule agonists of the TrkB receptor (Shibata et al., 2007; Yu et al., 2012; Yu et al., 2013), for the protection or regeneration of cochlear neural function caused by synaptopathic noise exposure. Using parallel *in vitro* and *in vivo* approaches, we characterized protection or rescue from cochlear synapse loss/deafferentation in the mouse cochlea. We found that TrkB agonists were strikingly effective in protection and restoration of synapses and neural function in noise-exposed ears.

## Materials and Methods

### Animals

Mice (CBA/CaJ) were born and raised in our acoustically monitored animal care facility (Sergeyenko et al., 2013) from breeders purchased from Jackson Laboratory (Bar Harbor, Maine). All experiments were conducted in accordance with the Public Health Service policy on Humane Care and Use of Laboratory Animals.

### *In vitro* co-culture of neonatal SGNs and de-afferented organ of Corti

*In vitro* co-culture of SGN and de-afferented organ of Corti was performed according to the method described previously (Parker et al., 2010; Tong et al., 2013). Briefly, isolated SGNs were treated with 0.25 % trypsin for 15 min at 37 °C. After neutralization of the trypsin with 10% FBS in DMEM, neurons were collected by centrifugation, triturated to a single-cell suspension and co-cultured with the de-afferented organ of Corti. The organ was dissected by removal of the membranous labyrinth, the stria vascularis and spiral ligament. Reissner’s membrane and the tectorial membrane were removed, and the organ was cultured in a well on a cover glass coated with 1:1 poly-L-ornithine (0.01%; Sigma-Millipore, USA) and laminin (50 μg/ml; BD Biosciences, USA). Explants were cultured in N2 and B27-supplemented DMEM, 1% HEPES solution (Millipore Sigma, USA), 1:1000 ampicillin, 1:300 fungizone at 37° C, 6.5% CO2. Amitriptyline (AT; 0.5 μM; Sigma, USA) was added at the beginning of the co-culture and cultures were stopped after 6 days. The number of innervated inner hair cells at the end of the incubation was quantified (Brugeaud et al., 2014; Parker et al., 2010).

### *In vitro* glutamate toxicity

Organ of Corti explants from postnatal day (P)3 to P4 mice of both sexes, after 24 h in culture, were exposed to a solution of 0.4 mM kainic acid (Abcam, USA) diluted in culture medium. After a 1-h kainic acid treatment, the explants were washed 6 times with culture medium and maintained for 24 h in 80 μl of culture medium alone or containing AT (0.1 μM) or 7,8-dihydroxyflavone (DHF; 0.5 μM; Sigma, USA). The cultures were then fixed and prepared for immunofluorescence.

### Acoustic overexposure

Noise exposure (8-16 kHz octave-band noise, 100 dB SPL, 2 h) was conducted on awake, adult (16-wk) mice as described previously (Kujawa and Liberman, 2009). The noise was created by a waveform generator (model WGI; Tucker-Davis Technologies), filtered (8-16 kHz, >60 dB/octave slope; Frequency Devices), amplified (D-75 power amplifier; Crown Audio) and delivered (compression driver; JBL) through an exponential horn into a tabletop reverberant exposure chamber. Mice were placed in a subdivided cage suspended directly beneath the acoustic horn of the sound-delivery speaker. Prior to each exposure, the noise was calibrated to the target 100 dB SPL using a quarter-inch condenser microphone (variability <1 dB across subdivisions).

### Drugs and delivery

To optimize drug access to the cochlea, direct infusion of drug into the posterior semicircular canal was performed. For these experiments, awake animals were noise-exposed and held for 48 h. They were then anesthetized (ketamine 100 mg/kg and xylazine 10 mg/kg, IP). A small hole was made in the posterior semicircular canal and 500 nl of drug suspension (25 mM AT in artificial perilymph; or 5 mM DHF in 10% DMSO in artificial perilymph; or artificial perilymph without drug) was injected at a rate of 91 nl/min. At the end of the infusion, the needle was left in place for 5 min, after which the surgical defect was repaired. Animals were held for 2 wks before testing and processing.

To assess effects with less invasive, systemic delivery of drugs, we focused on AT and animals received intraperitoneal (IP) injections of AT in saline or an equal volume of the saline vehicle alone according to the following treatment groups: 1) AT (12.5, 25 or 50 mg/kg) or saline once daily for 5-9 days with noise exposure 6 h after the 3^rd^ day of treatment; 2) AT (50 mg/kg) or saline once, 6 h prior to noise exposure; 3) AT (50 mg/kg) or saline once, 3 d after noise exposure; 4) AT (50 mg/kg) or saline once daily for 9 days beginning 3 d after noise exposure. Groups of mice were evaluated 24 h after noise to assess acute losses, 2 wks after noise, when threshold recovery for this exposure is complete and synapse loss has stabilized (Kujawa and Liberman 2009) or were held for one year post exposure to examine long-term effects.

### Cochlear function tests

Distortion product otoacoustic emissions (DPOAEs) and auditory brainstem responses (ABRs) were recorded from anesthetized mice (ketamine 100 mg/kg and xylazine 10 mg/kg, IP) in a 37°C, acoustically- and electrically-shielded chamber. A small incision made in the cartilaginous portion of the external ear canal provided a clear view to evaluate the health of the tympanic membrane and optimized acoustic system placement. Physiologic responses were stimulated and responses were collected using a 24-bit National Instruments PXI-based system controlled by custom LabView-based software. The acoustic system consisted of two miniature dynamic earphones (CDMF15008-03A; CUI) and a condenser microphone (FG-23329-PO7; Knowles) coupled to a probe tube.

Outer hair cell (OHC)-based DPOAEs were recorded in response to two primary tones, f1 and f2, with f2 equal to the frequencies used in ABR testing, f2/ f1 = 1.2 and L2 = L1-10 dB. For each frequency pair, DPOAE responses at 2f1-f2 were recorded, averaged, and analyzed as a function of level (L2: below threshold to 80 dB SPL) from sound pressure measurements in the ear canal. Threshold was defined as the L2 stimulus level required to generate a DPOAE amplitude of −5 dB SPL.

Neural-based ABRs were measured in response to tone pips (5.6 to 45.2 kHz; 0.5 ms rise-fall, 5 ms duration, 30/s, alternating polarity) using subdermal electrodes placed at the vertex, ventrolateral to the pinna, and at the base of the tail (ground). Stimuli were delivered at subthreshold levels to 90 dB SPL in 5 dB steps. Amplified (10,000X) responses were filtered (0.3-3 kHz), digitized, and averaged for each of the frequency-level combinations (1024 samples/level). Threshold was identified via visual inspection of stored waveforms as the lowest stimulus level needed to elicit a repeatable wave I response.

### Immunohistochemistry and quantification of afferent synapses

Immediately following the physiologic testing, anesthetized mice were transcardially perfused with 4% paraformaldehyde in 0.1 M phosphate buffer with additional perfusion through the cochlear scalae. Cochleas were post fixed in 4% paraformaldehyde for 1 h then decalcified (0.12 M EDTA) for up to 48 h. Microdissected pieces were placed in a blocking buffer (PBS with 5% normal horse serum and 0.3% Triton X-100) for 1 h at room temperature followed by overnight incubation at 37°C in antibodies to the following: (1) C-terminal binding protein 2 (mouse anti-CtBP2; BD Biosciences, used at 1:200). (2) myosin-VIIa (rabbit anti-myosin-VIIa; Proteus Biosciences; used at 1:200), and (3) GluA2 (mouse anti-glutamate receptor 2; Millipore; used at 1:2000). Cochlear pieces were incubated in species appropriate secondary antibodies coupled to Alexa Fluors in the red, blue, and green channels for 2 h at 37°C.

Cochlear piece lengths, collected and compiled using ImageJ software, were used to create a cochlear frequency map to localize cochlear structures to specific frequency regions (Muller et al., 2005). Confocal z-stacks were obtained (Leica TCS SP5) using a high-resolution, glycerin-immersion objective (63x, 1.3 N.A.) and a 3.17x digital zoom, 1024 x 512 raster. Two adjacent stacks (0.25 µm step size) were imaged for each targeted frequency region, each image spanning 78 µm of cochlear length. Confocal image stacks were ported to image-processing software (AMIRA; VISAGE Imaging) where inner hair cells (IHCs) and synapses were quantified as previously described (Fernandez et al., 2015; Sergeyenko et al., 2013). Synaptic associations between presynaptic IHC ribbons and post-synaptic glutamate receptors (synapses) were determined with custom software that computed and displayed the x-y projections of the voxel space within 1 um of the center of each ribbon identified by Amira analysis (Fernandez et al., 2015).

Immunohistochemistry and quantification of regenerated afferent synapses in the *in vitro* assays were performed by the same methods (Tong et al., 2013). Following fixation and permeabilization for 1 h, cultures were incubated with primary antibodies, anti-CtBP2 (mouse monoclonal IgG1; BD Biosciences, USA, used at 1:1,000), anti-PSD-95 (mouse monoclonal IgG2a; NeuroMab, USA, used at 1:1,000), and anti-neurofilament (chicken antibody to neurofilament-H; Millipore, USA, used at 1:2,500) overnight at 4° C. After rinsing three times for 10 min with 0.01 M PBS, pH□7.4. They were incubated with secondary antibodies: cyanine-5-conjugated goat anti-mouse IgG1 (Caltad Laboratories), biotin-conjugated Fluor goat anti-mouse IgG2a (Caltad Laboratories), Alexa Fluor 568-Streptavidin (Molecular Probes) and Alexa Fluor 488 goat anti-chicken (Molecular Probes) for 2 h at room temperature. Newly generated afferent synapses were identified by triple labeling of CtBP2, PSD-95 and neurofilament.

### Statistics

Statistical analyses were conducted using Prism 7.0 software (GraphPad). For analyses on adult mice, a one-way or two-way analysis of variance (ANOVA) was used, followed by Bonferroni correction for multiple comparisons. For analyses on neonatal mice, a two-tail, unpaired Students’ *t*-test was used. For all analyses, an alpha of < 0.05 was considered significant.

## Results

### AT and DHF increased cochlear synaptic regeneration in vitro

To assess synapse formation in a culture system, in which SGNs were co-cultured with de-afferented organ of Corti (Martinez-Monedero et al., 2006; Tong et al., 2013), the co-cultures were treated with 0.5 μM AT or maintained as controls. Six days later, immunohistochemistry demonstrated that cochlear afferent synapses had regenerated in the cultures (**Fig. 1A** and **B**). Quantitative measurement indicated a significant increase in synaptic regeneration in the AT treated cultures (**Fig. 1C**, * p < 0.05).

**Figure 1.**
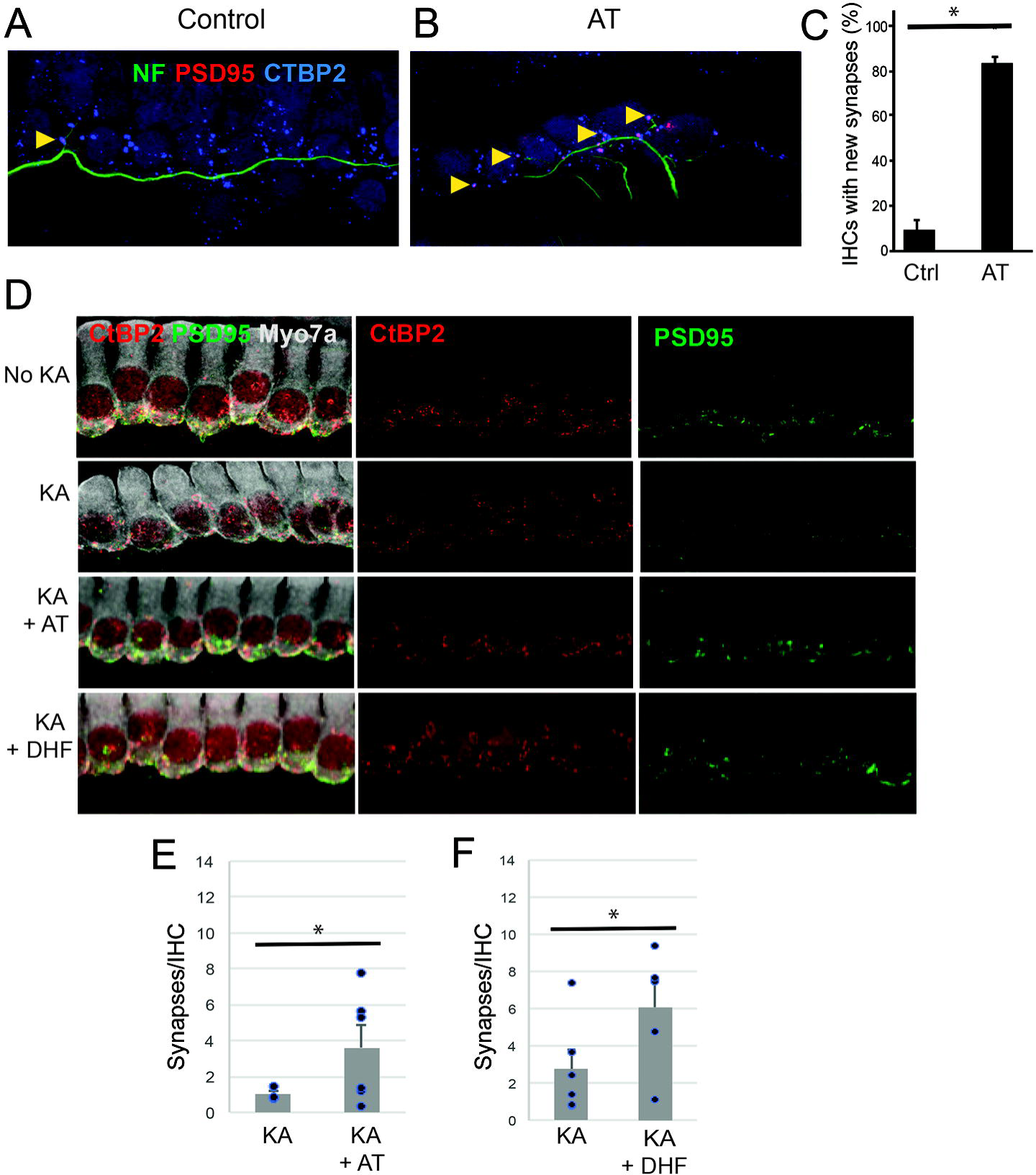
AT acts as a TrkB agonist in SGN. (**A-C)** Isolated SGNs and denervated organ of Corti were co-cultured in the absence (**A**), or presence (**B**) of 0.5 μM AT. After 6 days in culture, explants were fixed, immunostained with antibodies against neurofilament (green), CtBP2 (blue) and PSD-95 (red), and confocal images were obtained in the inner hair cell region (**A**,**B**). Juxtapositions of hair cell ribbons and afferent endings were identified by CtBP2/PSD-95 puncta (yellow arrowheads) and counted (**C**), indicating a significant increase in percentage of juxtaposed CtBP2/PSD-95 puncta at inner hair cells after AT treatment. Error bars indicate mean +/- SEM; asterisk indicates p<0.05. (**D-F)** Cochlear explants were cultured with (KA) or without (No KA) exposure to kainic acid (0.4 mM) for 1 h followed by treatment with culture medium with 0.1 μM AT (KA +AT) or 0.5 μM DHF (KA + DHF). The cultures were immunostained for CtBP2 (red) and PSD-95 (green) after 48 h (**D**). AT (**E**) and DHF (**F**) induced significant synaptic regeneration. Error bars indicates mean +/- SEM (n=3 for control and 6 for AT in **E**; n=6 for control and 6 for DHF in **F**); asterisks indicate p<0.05.

We adapted a rat model of excitatory cochlear synaptopathy (Wang and Green, 2011) to the mouse explants to assess the regenerative effect of the drug on cochlear synapses. In this model, cochlear explants, consisting of hair cells and attached SGNs, are exposed to selective ionotropic glutamate receptor agonist, kainic acid, to mimic glutamate toxicity, the excessive release of glutamate into the synaptic cleft with attendant damage to the terminal processes of SGNs (Pujol et al., 1985; Pujol and Puel, 1999). Synapses were detected by the occurrence of CtBP2-expressing synaptic ribbons and PSD95-positive post-synaptic densities. Damage was extensive after 1 h of treatment with kainic acid, and a small amount of reinnervation of hair cells by the peripheral processes of the neurons was apparent after 24 h in the absence of drugs (**Fig. 1D**). Addition of 0.1 μM AT (**Fig. 1E**) or 0.5 μM DHF (**Fig. 1F**) to the *in vitro* synaptogenesis assay resulted in a significant increase in juxtaposed ribbons and PSD95-positive neural endings relative to untreated samples.

### AT rescued synapses and auditory function *in vivo*

Prior studies in adult CBA/CaJ mice have demonstrated that exposure to a 2 h, 100 dB SPL octave band noise (8-16 kHz) results in a large, but reversible threshold shift, as evidenced by DPOAE and ABR wave-I measures. The exposure causes no acute loss of inner or outer hair cells but initiates an immediate and largely permanent loss of synapses between auditory nerve terminals and cochlear IHCs in mid to high frequency locations along the basilar membrane. This is accompanied by persistent declines in suprathreshold amplitudes of neural, but not OHC responses (Fernandez et al., 2015; Kujawa and Liberman, 2009, 2015). Here, we aimed to determine whether Trk agonist treatment could ameliorate some of the irreversible damage and reduce neural function caused by this noise exposure.

We reasoned that direct administration of the TrkB agonists to perilymph would be the best route for administration of the drugs *in vivo*. We delivered AT or DHF or vehicle alone into perilymph via posterior semicircular canal injection 48 h after noise exposure. When tested 24 h after exposure (before drug treatment), animals displayed similar ABR wave 1 threshold elevations at test frequencies above the noise band (21-45 kHz, Fig. **2A**). Thresholds recovered to within about 5 dB by 2 wks (**Fig. 2B**). Results are consistent with our previous reports (Kujawa and Liberman 2009; Fernandez et al., 2015) and demonstrate, further, that the drug treatments did not alter threshold recovery after noise.

**Figure 2.**
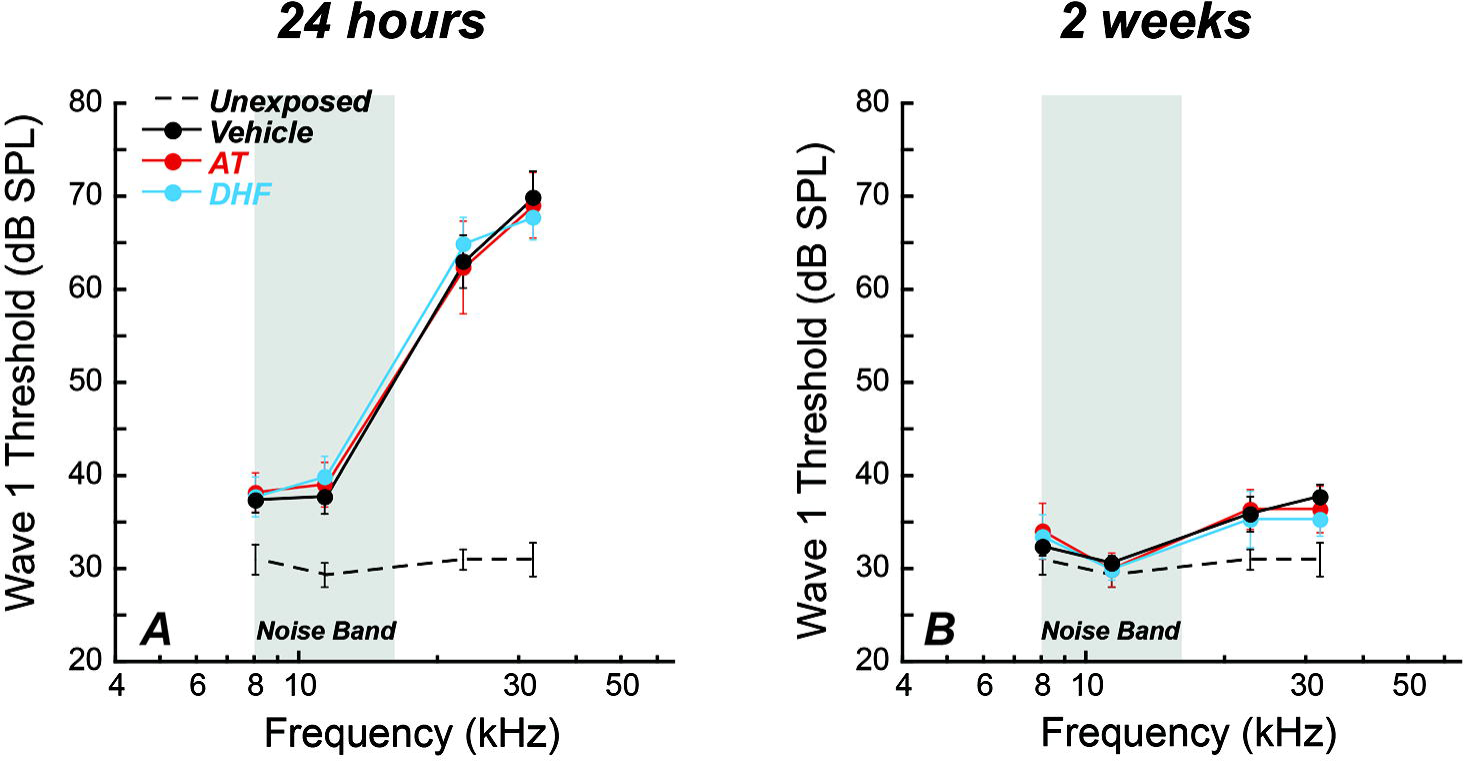
Noise-induced threshold elevations were similar across groups and Trk agonists did not alter recovery. **A:** Wave 1 thresholds recorded 24 h after noise (8-16 kHz, 100 dB SPL, 2h; gray bar) and before drug delivery to the posterior semicircular canal were similar across treatment groups (AT, 25mM; DHF, 5mM; AP-vehicle). Compared to strain- and age-matched, unexposed controls (Fernandez et al., 2015), maximum shifts were ∼40 dB at basal cochlear frequencies. **B:** By 2 wk post noise, all groups recovered to within ∼5dB of baseline values. Data shown as means ± SE, n=6 to 8/group).

Both AT- and DHF-treated mice demonstrated significantly greater neural response amplitudes than mice treated with the vehicle alone when assessed at the 2 wk post-exposure time point (2-way ANOVA, F_(2, 130)_=4.636, p=0.0114). In basal cochlear regions of maximum noise injury, vehicle-treated mice showed ∼45% wave 1 amplitude reductions compared to unexposed mice, whereas amplitudes recorded in AT- and DHF-treated animals recovered to ∼70% of unexposed controls (see for example 30 kHz, **Fig. 3A**). Proportionately, and in the same cochlear region, AT- and DHF-treated mice demonstrated significantly greater synapse counts than mice treated with vehicle alone (2-way ANOVA, F_(2,72)_=8.182, p=0.0006). Two weeks following exposure, synaptic declines of almost 50% were observed in the noise-damaged basal cochlea of vehicle-treated mice, whereas counts recorded in AT- and DHF-treated animals recovered to 70-75% of unexposed controls.

**Figure 3:**
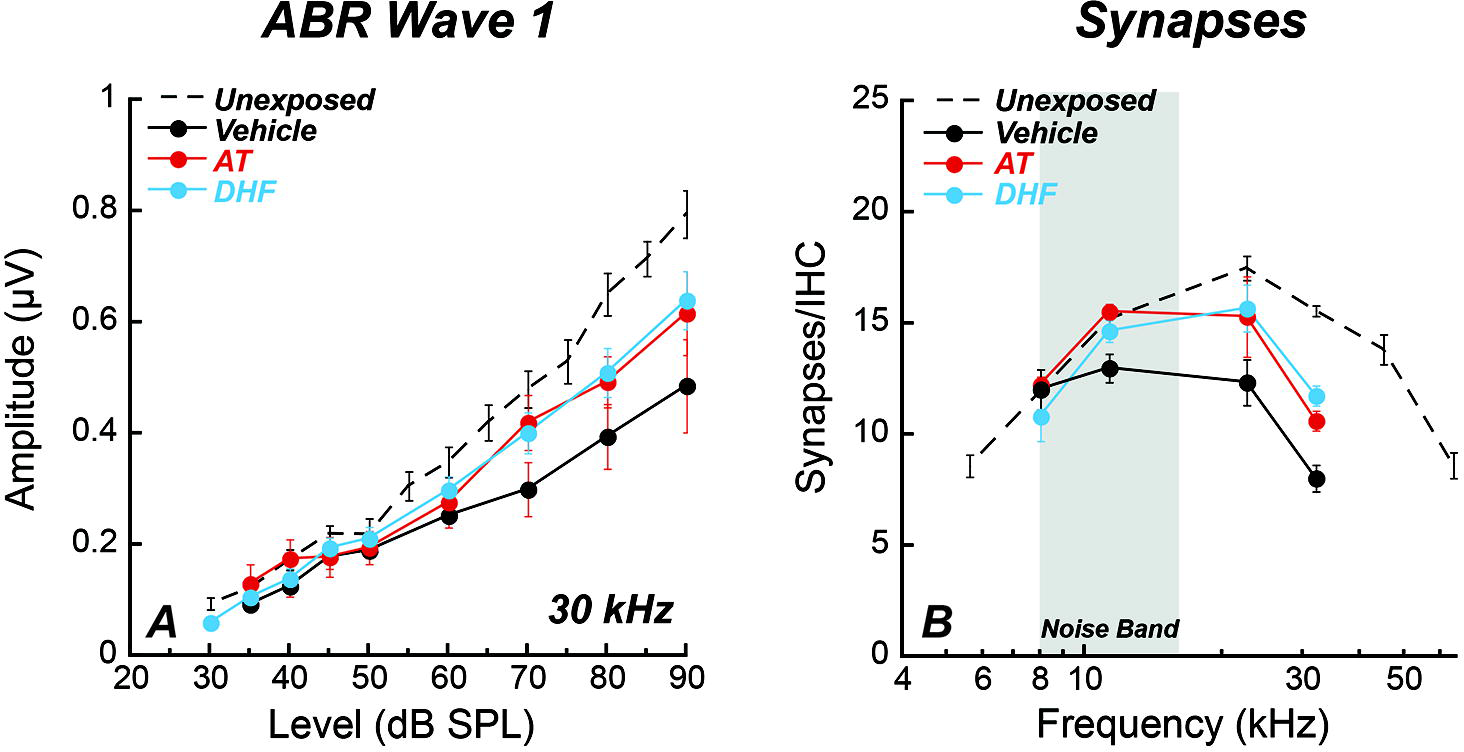
Local, post-exposure treatment with Trk agonists recovers cochlear nerve function and IHC synapses. Animals treated with 25 mM AT or 5 mM DHF 48 h after noise showed significantly greater ABR wave 1 amplitudes (F_(2, 130)_=4.636, p=0.0114) (**A**; shown for example at 30 kHz) and synapse counts (F_(2,72)_=8.182, p=0.0006) (**B**) compared to those treated with vehicle alone. Data shown as means ± SE, n=6 to 8/group. Unexposed controls from Fernandez et al., 2015. 2-way ANOVA with Bonferroni multiple comparisons test, *p<0.05, **p<0.01.

### AT preserved synapses and auditory function in vivo when administered systemically prior to noise-exposure

Given the effects of directly administered drug, and the potential to give these drugs systemically due to their low risk safety profile, we next asked whether the same effects could be achieved by systemic delivery. We performed these experiments with AT.

In our initial experiments, mice were treated systemically with saline or AT (12.5, 25, or 50 mg/kg) before and after noise exposure. Subsets of animals were tested 24 h after exposure and displayed expected and similar threshold elevations for both DPOAEs and ABR wave I at test frequencies above the noise band. Thresholds recovered by 2 wks as shown previously for this exposure (Kujawa and Liberman 2009; Fernandez et al., 2015; Fernandez et al., 2020). DPOAE amplitudes at 2 wks showed essentially complete post-exposure recovery for all drug and saline treatment groups (**Fig. 4a**). Together with the DPOAE and ABR wave 1 threshold recovery, the finding is consistent with functionally intact OHCs and no effect of the drugs on their post-exposure recovery.

**Figure 4.**
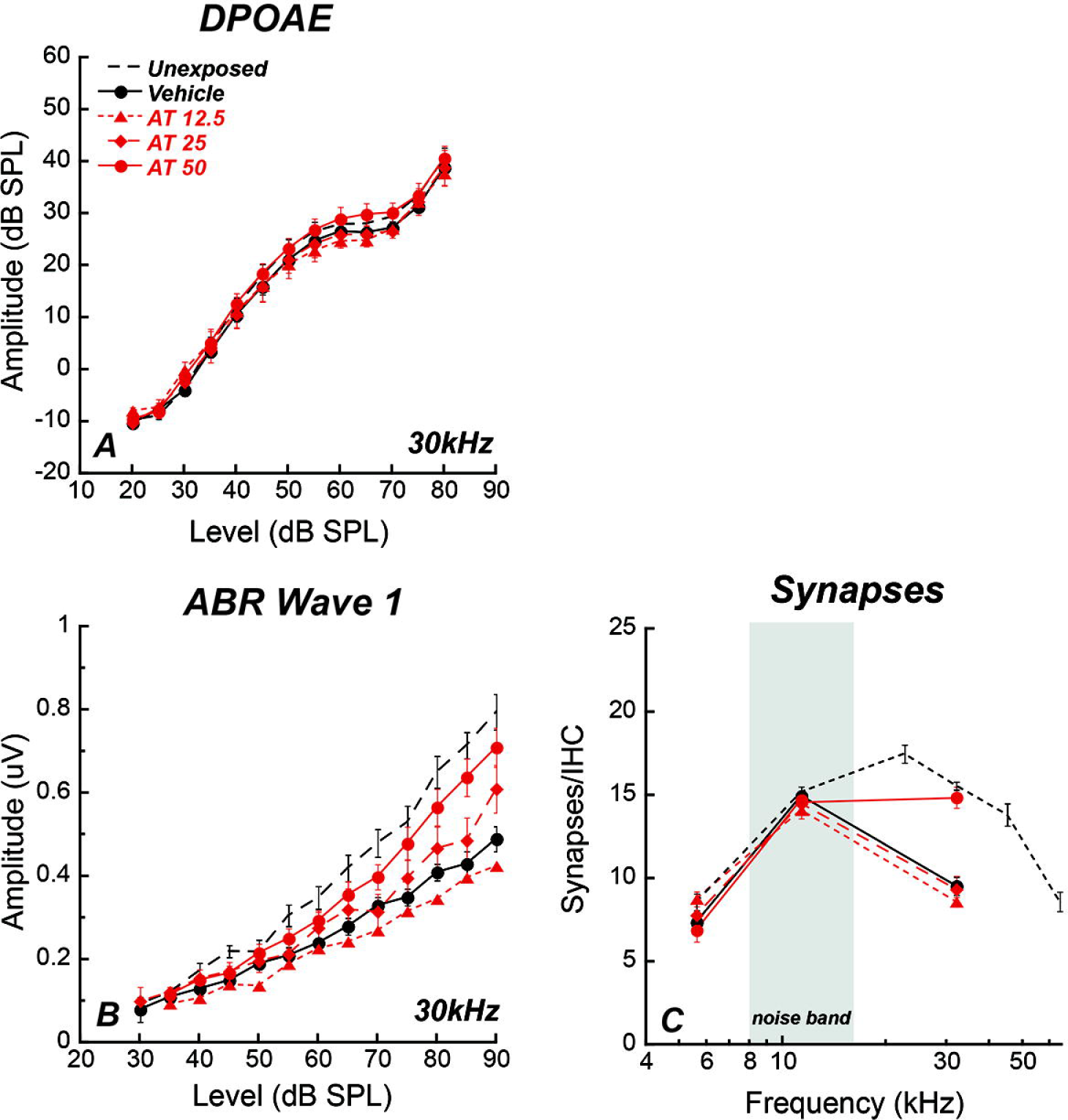
Systemic AT protection is dose-responsive. Amitriptyline (AT) is delivered systemically (12.5, 25 or 50 mg/kg once daily for 5-9 days). Animals are noise exposed on day 3, 6 hr after drug. In all groups, DPOAE and ABR thresholds and DPOAE amplitudes recover by 2 wks, but ABR amplitudes do not (**A**,**B**). Compared to saline vehicle controls, a significant preservation of ABR wave-I amplitude (**C**; F_(1,392)_=70.39, p<0.0001) and synapse counts (**D**; F_(1,116)_=18.90, p<0.0001) at 30 kHz is apparent in mice treated with 50 mg/kg AT. Data shown as means ± SE, n=10 to 30/group). Unexposed controls from Fernandez et al., 2015. 2-way ANOVA with Bonferroni multiple comparisons test, **p<0.01, ***p<0.001, ****p<0.0001.

In contrast, two weeks after noise, animals showed permanent ABR wave 1 amplitude decrements that varied with treatment and dose (**Fig. 4B**). At the highest dose of AT, 50 mg/kg, response amplitudes averaged over 80-80 dB were significantly larger [2-way ANOVA, F_(3,238)_=11.07, p<0.0001; Bonferroni multiple comparison significance at 17 kHz (p<0.03) and 30 kHz (p<0.001)] than those recorded for animals in the other groups. Synapse counts at cochlear regions spanning the frequency regions assessed physiologically also were greater in drug-treated animals (2-way ANOVA, F_(1,116)_=18.90, p<0.0001) (**Fig. 4C**). Vehicle-treated mice showed a maximum 50% synapse loss relative to unexposed controls, whereas they were nearly normal for AT 50 mg/kg-treated mice.

Subsequent experiments focused on the effective dose (50 mg/kg) and compared outcomes for treatments given only before or only after noise, as follows: 1) once, 6 h before noise, 2) once, 3 d after noise and 3) once daily for 9 days beginning 3 days after exposure.

Post-exposure threshold recovery by 2 wks was again unaltered by the treatments. However, wave 1 amplitudes and synapse counts at basal cochlear frequencies were sensitive to the timing of treatment relative to noise exposure; both were near normal when AT was given before (or before and after) noise, but declined to ∼50% when AT was administered only after noise (**Figs. 5A, B).** For animals receiving single dose pre-noise AT, suprathreshold wave 1 amplitudes and IHC synapse counts at 2 wks were ∼50-60% larger than saline controls in the noise-damage region (17 and 30 kHz). However, when the same AT dose was delivered 6 h after noise, neither ABR wave-I amplitudes (**Fig. 5A**) nor synapse counts (**Fig. 5B**) were systematically different from those recorded for saline controls: Amplitudes: F_(1, 58)_=3.742, p=0.0579; Synapses: F_(1, 112)_=0.06407, p=0.8006. Thus, AT showed potent protection but only at the highest dose (50 mg/kg), and a single injection at high concentration was as effective as repeated injections.

**Figure 5.**
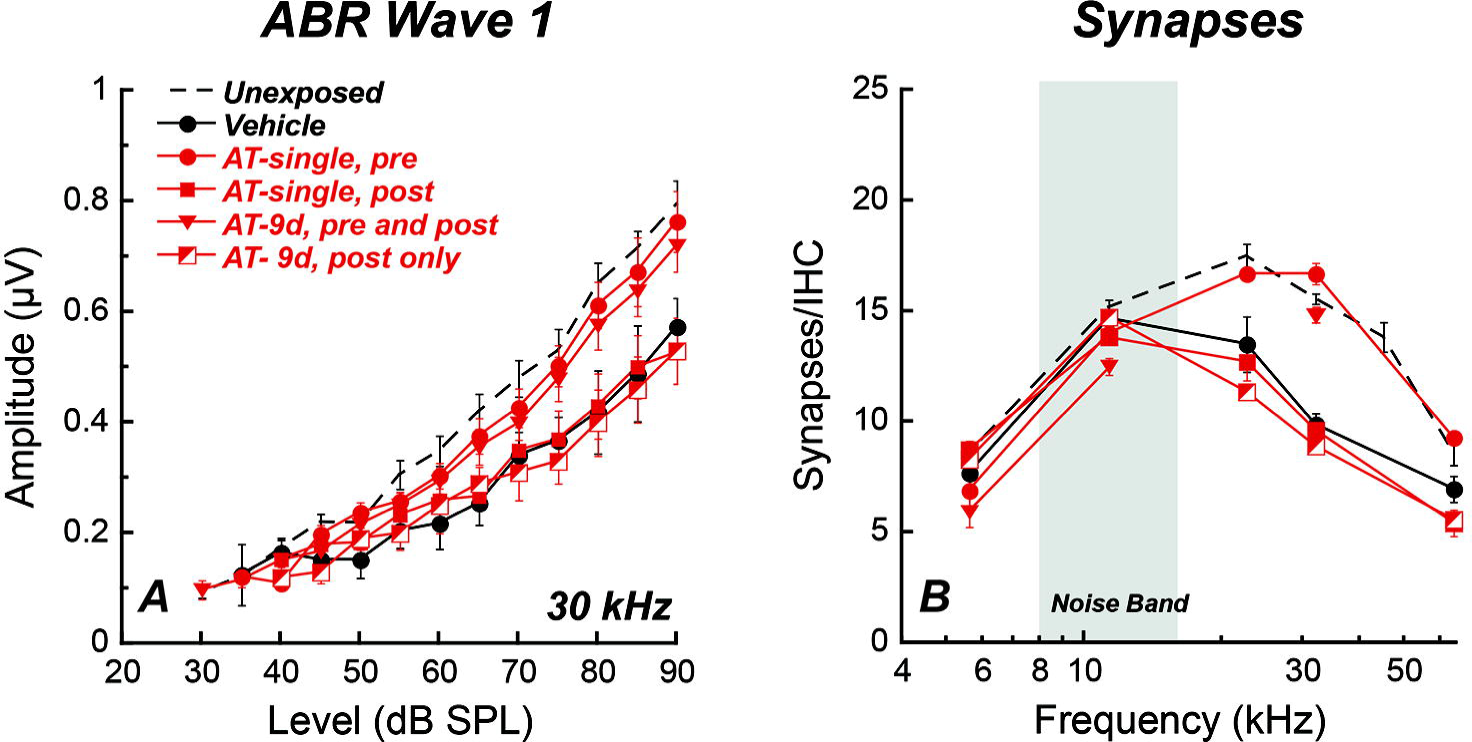
Systemic AT, delivered pre-, but not post-exposure protects cochlear nerve function and IHC synapses. Groups of animals received IP injections of AT in saline or saline vehicle alone according to the following treatment groups: 1) AT (12.5, 25 or 50 mg/kg) or saline once daily for 5-9 days with noise exposure 6 h after the 3^rd^ day of treatment; 2) AT (50 mg/kg) or saline once, 6 h prior to noise exposure; 3) AT (50 mg/kg) or saline once, 3 d after noise exposure; 4) AT (50 mg/kg) or saline once daily for 9 days beginning 3 days after noise exposure. Together, pre-exposure AT delivery was required to achieve protection (ABR Amplitude: F_(2,496)_=55.49, p<0.0001; Synapses: F(2,143)=24.19,p<0.0001). Data shown as means ± SE, n=10 to 30/group). Unexposed controls from Fernandez et al., 2015. 2-way ANOVA with Bonferroni multiple comparisons test.

### AT preserved synapses and auditory function in vivo one-year after noise exposure

Prior work has documented gradually-progressive loss of synapses with aging, and acceleration of these losses with spread to more apical regions in animals receiving synaptopathic noise exposure as young adults (Sergeyenko et al., 2013; Fernandez et al., 2015). Here, subsets of animals from the initial experimental series (systemic AT 12, 25, 50 mg/kg) were followed to 1 year post exposure to assess long-term effects of drug treatments on cochlear deafferentation. Shown in **Figures 6A** and **B** are DPOAE and ABR wave 1 amplitudes (30 kHz) for these long-held groups. OHC-based DPOAEs are similar across treated groups, documenting minimal exacerbation of ongoing declines relative to age-only mice (Fernandez et al., 2015). Of note, the neuroprotective treatment effect of AT apparent at 2 wks (**Fig. 4C**) persists, with mice that received pre-noise AT (50mg/kg) displaying significantly greater neural response amplitudes at 17 kHz (p<0.01) and 30 kHz (p<0.05) than vehicle-treated animals at the 1 yr test time (2-way ANOVA, F_(1,69)_=11.46, p<0.01 with Bonferroni multiple comparisons test). Indeed, with responses little different from never-exposed controls, these animals appear to have been fully protected from the noise-induced cochlear de-afferentation.

**Figure 6:**
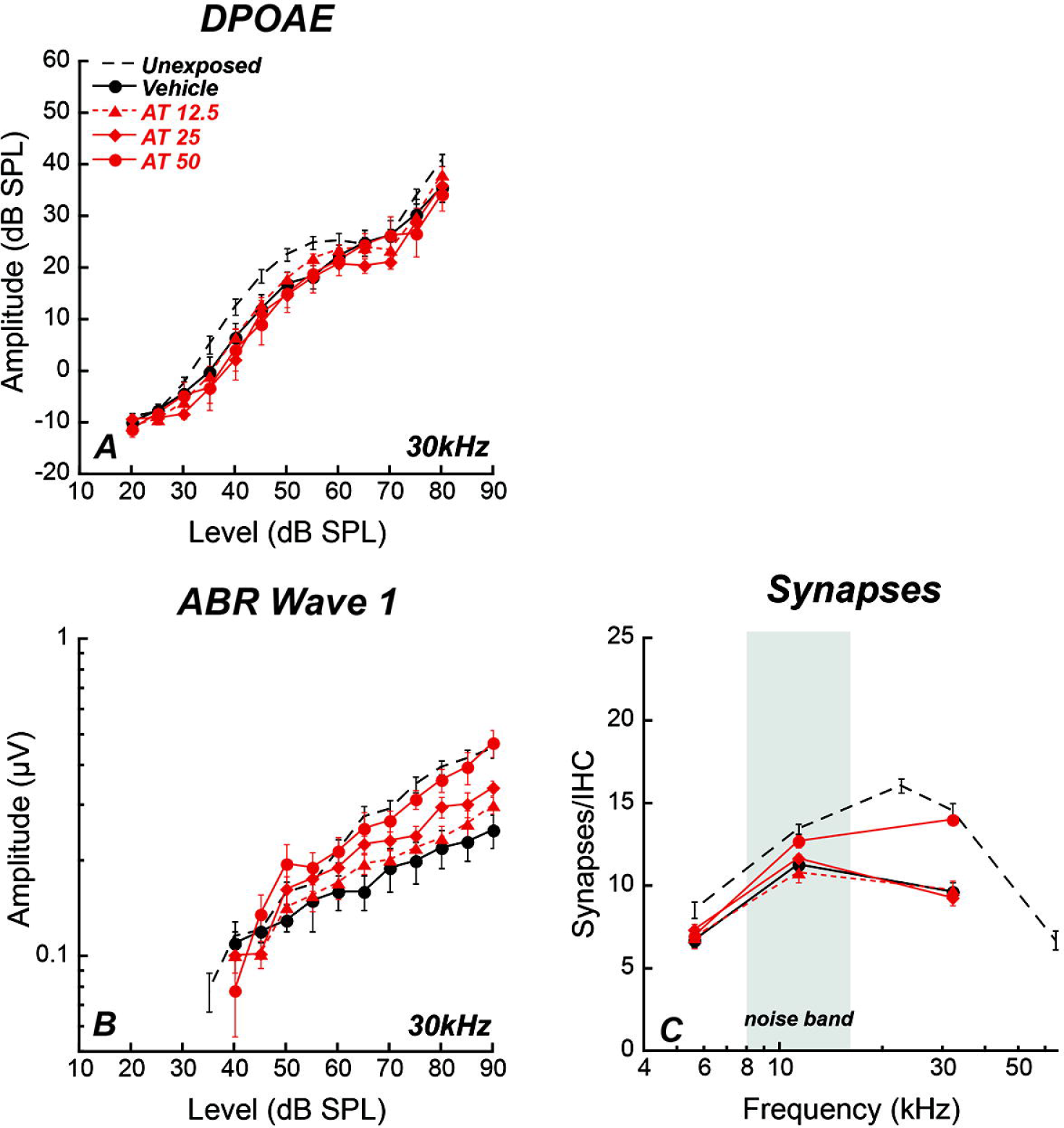
AT protection is long-lasting. Subsets of mice were held 1 year post noise. DPOAE amplitudes remained well preserved in all groups (**A**). Relative to age-matched, vehicle treated controls, significant conservation of ABR wave I amplitude (B) was observed in mice treated with AT 50 mg/kg for 9 days beginning 3 days pre noise (F_(1,122)_=33.88, p<0.0001) (**B**). The number of synapses remaining at 2 wks also was also maintained 1 year after noise exposure (**C**), with significantly more synapses in the high frequency 32 kHz region for mice treated with AT (F_(1,131)_=16.71, p<0.0001). Group means ± SEMs are shown: n= 7-30 mice per group. Unexposed controls from Fernandez et al., 2015. 2-way ANOVA with Bonferroni multiple comparisons test, *p<0.05, **p<0.01, ***p<0.001, ****p<0.0001.

Support for this notion also can be found in the synapse counts from these mice (**Fig. 6C**). High-dose AT treated mice showed significantly greater (F_(1,131)_=16.71, p<0.0001) synapse counts than vehicle-treated ears at 30 kHz (p<0.0001) where counts were consistent with age-matched, unexposed mice. Moreover, as documented previously, the damage region in aging vehicle-treated ears spread apically toward the previously unaffected region of 11k Hz, where synapse counts were now 20% greater in age-matched, AT-treated mice.

When compared to data collected for animals held 2 wks following noise exposure, wave 1 amplitude differences between saline and AT-treated mice were exaggerated at 1 yr, with vehicle-treated animals showing greater ongoing declines after noise. Of note, DPOAE amplitudes recorded from the same ears were comparable for all groups. Together, findings show that drug-related effects target synapses/neurons rather than OHCs, effects are long-lasting, and the increasing separation between saline and AT (50 mg/kg) is not reflected at the level of the OHCs. Additionally, as synapse counts in saline-treated animals show results consistent with our previous reports in exposed, then aged mice (Fernandez et al., 2015), the significantly smaller age-progressive declines for AT-treated mice suggest that the drug also may have long-term protective effect against age-progressive de-afferentation.

## Discussion

We have shown here that treatment with AT restores synapses between cochlear afferent neurons and sensory inner hair cells. In contrast to the ears of control (saline-treated, noise-exposed) animals, in which extensive noise-induced afferent synapse loss persists after thresholds recover, synapses were significantly protected by pre-exposure treatment and rescued by post-exposure treatment with AT, and cochlear neurons appeared functionally intact. The protective effect of pre-noise AT against cochlear deafferentation was long-lasting. The regenerative effect of the drug is particularly significant for clinical application, as exposure to noise, which may be a principal contributor to synaptopathy in the human population, can occur in varied environments with unpredictable timing and may be cumulative. The effect of AT in mitigating the synaptic loss and promoting synaptogenesis was also seen *in vitro*, providing support for a regenerative mechanism and pointing to TrkB receptor involvement.

Synaptopathy occurs as a primary consequence of noise exposure and can permanently reduce cochlear afferent response amplitudes in ears with functionally intact hair cells and normal thresholds (Kujawa and Liberman, 2009). Correlation between loss of amplitude and synaptic loss suggested a ‘hidden hearing loss’ in which audiograms (assays of threshold sensitivity) could recover to normal and the loss of synapses preceded the loss of hair cells and could act independently of it. However, even when thresholds recover, this cochlear deafferentation may compromise the fidelity of suprathreshold signal coding and moreover appears to be a harbinger of exaggerated declines in hearing function (Kujawa and Liberman, 2006, 2009). Indeed this type of mechanism for loss of neural response amplitude has now been shown to be important in the loss of cochlear function caused by aging as well as noise exposure (Sergeyenko et al., 2013).

The activity we find here for a Trk agonist is consistent with previous data on Trk receptor activation by Trk ligands, NT3 and BDNF (Ramekers et al., 2015; Ruel et al., 2007; Shepherd et al., 2005; Shinohara et al., 2002; Wise et al., 2005). NT3 and BDNF have long been shown to be protective or ameliorative for neural loss in the cochlea. The roles of NT3 and BDNF may be different in synaptogenesis, fiber growth, and survival. The latter roles, neuron survival and growth, are the most thoroughly characterized of the effects and are found throughout the nervous system. NT3 and BDNF do not appear to be equivalent for these roles (Fritzsch et al., 2006; Green et al., 2012; Tessarollo et al., 2004). Both neurotrophins are present in the inner ear, and several studies have shown that NT3 is the principal neurotrophin in the cochlea, whereas BDNF has a more important role in the vestibular system (Fritzsch et al., 1997). However, there is considerable overlap of function. Indeed, the two neurotrophins have been used together in many studies (Wise et al., 2005), and both stimulate neurite outgrowth and neuron survival.

Neurotrophins also show distinct actions in regeneration of neurons. This distinction may be key to the effects of AT on regeneration vs protection. AT has been shown to have TrkB activity with no TrkC activity (Jang et al., 2009). There are some indications that NT3 in particular is effective in cochlear hearing loss, whereas BDNF is effective in vestibular dysfunction due to synaptic lesions (Gomez-Casati et al., 2010; Suzuki et al., 2016; Wan et al., 2014). NT3 but not BDNF appeared to reverse the synaptopathy in a genetic model (Wan et al., 2014), and NT3 application to the round window (Suzuki et al., 2016) protected synapses from noise induced synaptopathy. If indeed NT3 stimulation of TrkC, but not BDNF stimulation of TrkB can repair cochlear synapses, then it seems logical that a TrkB agonist would be suboptimal. As we see a protective effect when the drug is given prior to noise exposure, as well as a regenerative effect when the drug is given after noise exposure, we believe that TrkB stimulation is effective for the stimulation of synaptogenesis in the cochlea. The effect of AT suggests that TrkB agonist activity is sufficient to restore synapses. It is hard to compare to the results on NT3 vs BDNF in transgenics where the concentration of peptide is unknown and may differ from the BDNF equivalents we added. However, we cannot rule out the possibility that high concentrations of the drug could also produce some activation of TrkC.

BDNF has a stimulatory effect on regeneration in many systems and is released in response to damage. In studies of aminoglycoside induced hearing loss, ears that were administered neurotrophins after the insult showed less neural loss (Ramekers et al., 2015; Shinohara et al., 2002; Wise et al., 2005); however, this effect was ascribed to protection from neural loss that occurs secondary to the loss of hair cells. Regeneration of synapses was reported in the newborn rat *in vitro* (Wang and Green, 2011). In that case the synapses partially recovered spontaneously, and the recovery could be augmented by BDNF or NT3. However, the endogenous factor responsible for the spontaneous regeneration appeared to be NT3. Protection of hearing from noise exposure was correlated to secretion of BDNF that occurred at night but not during the day (Meltser et al., 2014). In the studies reported here, administration of AT after the noise exposure restored synapses when it was at sufficient concentrations. Posterior semicircular canal delivery proved the most effective route and was presumed to be successful because drug reached the cochlear perilymph at sufficient concentrations. Given the regenerative effect of AT on cochlear neurons, the possibility remains that the observed protective effect was also a regenerative effect occurring after the noise exposure, i.e. a fast regeneration comprising synaptic remodeling on a rapid time scale as occurs in synaptic spines in a process driven by BDNF (Vignoli et al., 2016). The time window we looked at for administration of the drug was limited to the period shortly (2 days) after noise exposure. Development of a regenerative treatment using this drug will require assessment of the full extent of the post-damage window for a therapeutic effect.

Tricyclic anti-depressants like AT have Trk agonist activity (Jang et al., 2009). Interestingly, this activity was discovered long after their introduction for the treatment of mood disorders. The compounds were initially used for inhibition of serotonin and noradrenaline transport (Snyder and Peroutka, 1982; Snyder and Yamamura, 1977; U’Prichard et al., 1978). Since the discovery of the neurotrophic activity the compounds have been tested for the amelioration of various types of hearing loss (Shibata et al., 2007; Yu et al., 2012; Yu et al., 2013). The activity on a genetic hearing disorder was striking (Yu et al., 2013) and was thought to mimic BDNF activity. The effect on noise damage was thought to be due to protection of SGNs (Shibata et al., 2007). AT is a well-tolerated drug that has been tested thoroughly for mood disorders, and our data suggest that AT or related drugs could have important clinical applications in noise-induced hidden hearing loss as studied here and potentially in age-related hidden hearing loss related to cochlear deafferentation.

## Notes

### Competing Interest Statement

The authors have declared no competing interest.

